# Operationalizing ecological connectivity in spatial conservation planning with Marxan Connect

**DOI:** 10.1101/315424

**Authors:** Rémi M. Daigle, Anna Metaxas, Arieanna Balbar, Jennifer McGowan, Eric A. Treml, Caitlin D. Kuempel, Hugh P. Possingham, Maria Beger

## Abstract

1. Globally, protected areas are being established to protect biodiversity and to promote ecosystem resilience. The typical spatial conservation planning process leading to the creation of these protected areas focuses on representation and replication of ecological features, often using decision support systems such as Marxan. Unfortunately, Marxan currently requires manual input or specialised scripts to explicitly consider ecological connectivity, a property critical to metapopulation persistence and resilience.
2. “Marxan Connect” is a new open source, open access Graphical User Interface (GUI) designed to assist conservation planners in the systematic operationalization of ecological connectivity in protected area network planning.
3. Marxan Connect is able to incorporate estimates of demographic connectivity (*e.g.* derived from tracking data, dispersal models, or genetics) or structural landscape connectivity (*e.g.* isolation by resistance). This is accomplished by calculating metapopulation-relevant connectivity metrics (*e.g.* eigenvector centrality) and treating those as conservation features, or using the connectivity data as a spatial dependency amongst sites to be included in the prioritization process.
4. Marxan Connect allows a wide group of users to incorporate directional ecological connectivity into conservation plans. The least-cost conservation solutions provided by Marxan Connect, combined with ecologically relevant post-hoc testing, are more likely to support persistent and resilient metapopulations (*e.g.* fish stocks) and provide better protection for biodiversity than if connectivity is ignored.

## Introduction

Connectivity, in its most general form, refers to the exchange of individuals (including genes, traits, disease, etc.), energy or materials among habitat patches, populations, communities or ecosystems. Maintaining connectivity can improve population resilience to perturbations, increase metapopulation viability, promote genetic diversity and maintain energetic pathways among ecosystems (Palumbi 2003; Figueira & Crowder 2006; Lowe & Allendorf 2010). Connectivity also appears at the forefront of global international conservation policy, so as Aichi Target 11,which commits 197 countries to establishing “effective, representative, and well-connected” networks of reserves by 2020 (UNEP 2010).

There are many metrics and methods to evaluate the connectivity of sea/landscapes and these can be used to assess networks of protected areas and influence spatial conservation planning in the future (Saura & Pascual-Hortal 2007; Beger *et al.* 2010a; Chollett *et al.* 2017; D’Aloia *et al.* 2017; Zeller *et al.* 2018). The quantity and quality of empirical data used to calculate connectivity have been growing rapidly in the last few years (Kool, Moilanen & Treml 2013; Hussey *et al.* 2015; Magris *et al.* 2018; Zeller *et al.* 2018). In turn, methods for estimating ecological connectivity are also advancing, and new conservation planning tools are quickly emerging to capitalize on these new data and methods (Saura & Pascual-Hortal 2007; Beger *et al.* 2010b; White *et al.* 2014). Examples of connectivity data that have been incorporated in conservation applications include: gene flow (Beger *et al.* 2014; Marrotte *et al.* 2017), dynamic distributions and migratory bottlenecks on migratory pathways (Iwamura *et al.* 2013; Runge *et al.* 2016), maximizing larval flow (Magris *et al.* 2016; D’Aloia *et al.* 2017), ontogenetic shifts in habitat use (Brown *et al.* 2016; Weeks 2017), ensuring the movement of adult individuals pathways (Beger *et al.* 2015; Mazor *et al.* 2016; Pereira, Saura & Jordán 2017; Zeller *et al.* 2018), and maintaining fisheries benefits (Daigle, Monaco & Elgin 2017; Krueck *et al.* 2017). Despite these efforts, connectivity is not commonly being incorporated in on-the-ground decision making for planning (Beger *et al.* 2010a; Barnes *et al.* 2018; Balbar, unpublished data). This is largely because connectivity metrics are not well defined or standardized, practitioners often lack confidence in the data or the expertise to work with them, and approaches to explicitly incorporate connectivity patterns in spatial planning are rare.

Spatial conservation planning is an approach that guides the allocation of conservation resources to areas identified as important for biodiversity whilst minimising the conservation impact on resource users (Margules & Pressey 2000; Moilanen, Wilson & Possingham 2009; Wilson, Cabeza & Klein 2009). The process of spatial planning demands setting broad goals, which can be turned into quantifiable objectives that lead to the conservation of biodiversity (Tear *et al.* 2005) and which, in turn, link back to actions, costs and feasibility (Wilson *et al.* 2007). Spatial planning often relies on the use of decision-support software (*e.g.* Marxan or Zonation) to help decide the location and timing of actions (*e.g.* establishing protected areas) to best achieve conservation objectives. These tools are primarily used to develop representative and cost-efficient conservation plans by meeting targets for species or habitats, with the consideration of connectivity patterns primarily expressed by prioritising adjacent or contiguous sites. To advance the inclusion of ecological connectivity into the spatial planning process, technical documentation, best-practice guidelines and user-friendly tools are needed. Knowing how to best identify, evaluate, and treat connectivity data to meet different objectives within a given spatial planning framework is important to better capture key ecological processes in planning.

Here, we outline potential workflows of realising connectivity in spatial planning, including the treatment of various data formats, key decision points that link back to objectives, types of data related to connectivity, evaluation and post-hoc analysis. We do so in the context of the widely used spatial planning tool Marxan, which aims to represent biodiversity whilst minimizing overall cost (Ball, Possingham & Watts 2009). We then introduce a new open source and open access tool called Marxan Connect to help users operationalize these concepts within Marxan. Our objective is to enable an overview of the selection and treatment of connectivity data to encourage its use in spatial conservation planning.

#### Box 1

A primer for spatial conservation planning with Marxan Marxan uses a simulated annealing algorithm to find good solutions to the “minimum set” problem. In the minimum set problem, the user specifies an amount of each conservation feature *j* that needs to be conserved, or conservation targets (*T_j_*), for each conservation feature. The basic minimum set problem is an integer linear programming problem and does not consider connectivity:

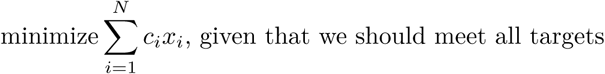

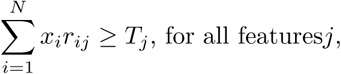

where *N* is the number of planning units, *c_i_* is the cost of planning unit *i*, *r_ij_* is the amount of feature *j* in planning unit *i*, and *x_i_* is a control variable which has the value of 1 for selected sites and 0 for unselected sites. It is usually desirable to include some basic spatial properties of a protected area system such as geographic proximity or adjacency information between planning units to help minimize costs or maximize clumping of a protected area system. For example, if the common boundary between every pair of planning units is known, then the minimum set problems can be extended to include a term for the boundary length of the reserve system and an effort made to minimise it:

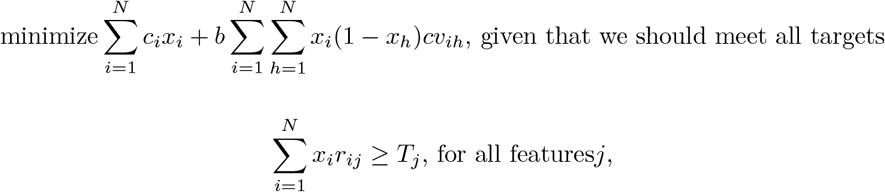

where *b* is the boundary length modifier (BLM), and *cv_ih_* represents the cost of a boundary and is typically the length of the physical boundary between sites *i* and *h*. Costs (*c_i_*) in Marxan often pertain to socio-economic implications of protecting a site, such as management or opportunity costs. For more information see Ball et al. (2009) and Ardron et al. (2010). Key terms and definitions:

- **Planning area**: the spatial domain over which the planning process occurs. This is synonymous with terms “domain” or “extent” or “study area” in other fields. This area is subdivided into smaller “Planning Units”.
- **Planning unit**: spatial units within the entire planning area (*i.e.* domain, or study area), which can be defined using regular gridded (*e.g.* hexagonal) or using landscape features-based (*e.g.* reefs, water catchments) as in Marxan.
- **Boundary Length**: the shared boundary length between adjacent planning units.
- **Boundary Length Modifier (BLM)**: a weighting parameter to ‘tune’ the influence of the boundaries. The BLM helps achieve “clumped” solutions by reducing the overall edge to area ratio. A higher BLM value results in a more ‘clumped’ Marxan solution.
- **Conservation feature**: the features (*e.g.* habitats, species, processes) for which a target is set.
- **Conservation target**: the minimum quantity or proportion of the conservation feature in the study area to be included in solutions.
- **Solution**: a binary output of Marxan reflecting whether a planning unit is selected (1) or not selected (0) as part of the conservation plan.
- **Selection Frequency**: the summed solution output of Marxan reflecting how many times a planning unit was selected across runs

## Understanding connectivity data

One of the challenges associated with integrating ecological connectivity in spatial planning is the wide variety of entities that move (*e.g.* organism, gene, pollutant) and movement processes (*e.g.* migration route, larval dispersal, multi-generational gene flow, carbon flux). While there are many types of data sources, connectivity data are often stored as matrices, where donor (or source) sites are rows, and the recipient (or destination) sites are columns. Alternatively, connectivity data may be stored in an edge list where the first column contains the donor site IDs, the second column contains the recipient site IDs, and the third column contains the connectivity value. Below, we review a few of the most common data sources organized by their treatment in Marxan Connect. Additional details on data format, types, mathematical representations and associated assumptions can be found on the Marxan Connect tool website, marxanconnect.ca.

### Landscape-based estimates of connectivity

Some spatial planners may have access to detailed connectivity information based on demographic data (See “Demographic estimates of connectivity” section below). In these cases, Marxan Connect can generate estimates of connectivity strength (*e.g.* spatial isolation) based either on the Euclidean distance between habitats, or isolation by resistance (McRae & Nürnberger 2006). These landscape-based connectivity estimates are often more limited in their applicability than demographic data (*e.g.* self-recruitment), but require less data.

#### Linkages across a habitat matrix

The structure and spatial configuration of the land- or sea-scape (*i.e.* habitat type, size, and spacing) can impede or facilitate the movement of organisms. The rate at which impediment or facilitation occurs has been defined as the strength of landscape connectivity (Tischendorf & Fahrig 2000). The impediment or facilitation (*i.e.* resistance or cost to traverse landscape) posed by habitat types can be estimated from tracking data, genetic data, expert opinion, or habitat suitability models for species-centric approaches (Bunn, Urban & Keitt 2000; Urban & Keitt 2001; Ricketts 2001; Zeller, McGarigal & Whiteley 2012). For a habitat or multi-species centric approach, resistance can also be estimated from the similarity in environmental variables (*e.g.* land cover) or that of species assemblages (Schumaker 1996). From this resistance surface, it is possible to estimate the rate of movement of organisms across the landscape based on the spatial arrangement of habitat patches using various methods such as least-cost path analysis, and current density approaches (Fall *et al.* 2007; Rayfield, Fortin & Fall 2010; Koen *et al.* 2014). While these methods are conceptually similar, they produce qualitatively different connectivity estimates and may be difficult to validate (Saura & Pascual-Hortal 2007; Rayfield, Fortin & Fall 2010; Zeller *et al.* 2018).

The conservation implications of these differences in connectivity have not been fully explored in the context of spatial planning. While methods based on Euclidean distance can be species non-specific, resistance-based models necessarily focus on specific species (Ricketts 2001). However, some features of the landscape (*i.e.* calculated using the multi-species approach) may have different, but important uses for multiple species. For example, current density is a metric adopted from electrical circuit theory which, in movement ecology, is intended to represent the prevalence of movement of organisms across a landscape. However, some species (amphibians and reptiles) may use areas of high current density as movement pathways while others (fishers - *Martes pennanti*) use these areas as home ranges (Koen *et al.* 2014).

### Demographic estimates of connectivity

Whether the movement of ‘individual’ organisms, particles of detritus or pollutants are being directly measured (tagging and tracking) or estimated (genetics or models), Marxan Connect treats these data as “demographic connectivity”(See marxanconnect.ca for more details on the mathematical representations of connectivity data). The strength of connectivity is measured as a probability or an absolute amount.

#### Tagging and Tracking

Movement ecology has seen profound advances over the last decades arising from the ability to identify and track individuals and thus understand the movement of organisms in space and time (Hussey *et al.* 2015). Traditionally, individual organisms have been identified using scarring, banding, tagging, radiotelemetry collars, passive integrated transponder devices (PIT tags), otolith/statolith microchemical signatures, or other approaches that allow an observer to track the movement of organisms through a landscape (Scott 1942; Thomas & Marburger 1964; Dunn & Gipson 1977; Twigg 1978; McNeil & Crossman 1979; Whitfield Gibbons & Andrews 2004). The advent of Global Positioning System (GPS) tags, satellite communication, and other forms of data relay provide opportunities to collect tracking data at higher spatiotemporal resolution and on a wider range of organisms, such as long-distance migratory species (Voegeli *et al.* 2001; Cagnacci *et al.* 2010).

Regardless of the approach, marking and tagging observations typically consist of a sequence of times and locations at which an individual was observed. The sequence often records numerous processes including ontogenetic movements, foraging, seasonal migrations, etc. The data can be used in spatial planning for objectives related to habitat use (*e.g.* identifying core foraging areas), movement pathways (*e.g.* finding migration routes) and species demography (*e.g.* disease spread through a population; McGowan *et al.* 2017b). Since tagging or tracking data are typically collected from relatively few individuals, they are often spatially biased and do not record multi-generational variation in movement.

#### Genetic approaches for estimating ecological connectivity

Genetic approaches have long been used to estimate the degree to which populations have diverged, and the degree to which gene flow via dispersal or migration influences this divergence (Palumbi 2003). As molecular techniques have changed, divergence and gene flow can be now be quantified at much finer spatial and temporal scales (Manel & Holderegger 2013; Hand *et al.* 2015). The integration of this fine-scale genetic information is being proposed as central to conservation and management efforts, despite technical and conceptual challenges (Beger *et al.* 2014). Generally, assignment methods are being used to develop estimates of ecological connectivity at the scale of populations (*e.g.*, using Structure Pritchard, Stephens & Donnelly 2000), whereby individuals are ‘assigned’ back to the population of origin. Similarly, parent-offspring assignments can be developed based on DNA fingerprinting to identify realised dispersal events over a single dispersal event (*e.g.* Saenz-Agudelo *et al.* 2009). In both cases, ecological, or demographically-significant, connectivity estimates between conservation planning units (and/or populations) can be estimated, if sampling is thorough and consistent with the planning unit structure. The result of these genetic approaches is often a migration matrix representing the likelihood that individuals or genotypes found at some destination populations came from the suite of sampled source populations. Unfortunately, these data are expensive to collect at the scale and scope appropriate for conservation applications, and appropriately interpreting the connectivity results is often highly context dependent. To date, the few published academic studies attempting this have struggled with significant compromises in taxonomic coverage, geographic extent, and alignment with the planning process (Harrison *et al.* 2012; *e.g.* Beger *et al.* 2014).

#### Individual-based models of movement or dispersal

Models of movement for individual organisms can be very useful in estimating connectivity. These models are typically based on the physical environment (*e.g.* ocean currents) and behavioural (*e.g.* resource selection) processes which influence movement. To model the movement of materials or individuals through land and seascapes, advection-diffusion models are often used as they can efficiently capture the physical fluid-like transport through these environments, including turbulence (Metaxas 2001), and other biological traits such as behaviour and duration of the dispersal phase (Roughgarden, Gaines & Possingham 1988; Cowen *et al.* 2000; Bode, Bode & Armsworth 2006; Paris, Chérubin & Cowen 2007; Metaxas & Saunders 2009; Treml *et al.* 2015; Daigle, Chassé & Metaxas 2016). In general, these modelling approaches are used to calculate the probability of exchange of individuals between habitat patches. Particle tracking approaches link source populations to destinations and vice versa allowing one to calculate the probability of linking any source to any destination and vice versa. These probabilities can then be used as the connectivity strength. Behaviour motivated movement requires some specific knowledge of the species being modelled. This behaviour can be based on resource availability (Chetkiewicz & Boyce 2009), predator avoidance (Bracis 2015), or other environmental cues (Daigle & Metaxas 2012; Zeller *et al.* 2018). While it is computationally feasible to scale up these models, it is often difficult to appropriately validate these complex models at the broad-scale most relevant to area-based management.

## Using connectivity data in spatial planning

In addition to the often used “rules of thumb” for connectivity which guide sizing and spacing of marine protected areas (Mora *et al.* 2006; Smith & Metaxas 2018), there are several different methods to directly incorporate connectivity data into spatial planning tools. For Marxan specifically, these include: 1) treating connectivity properties of planning units as conservation features (continuous or discrete) for which a target is set; 2) including connectivity strengths among planning units as spatial dependencies within the objective function; 3) treating connectivity properties of planning units as a connectivity cost; and 4) customizing the objective function to incorporate connectivity performance metrics. Methods 1) and 2) are fully implemented within Marxan Connect (Figure 2) while 3) is supported, but not facilitated for reasons described below. Method 4) is currently an area of active research not yet implemented. We note that in the following sections, our objective is to identify and discuss different treatments of connectivity data. We follow up with notes on making decisions related to data or methods, and we stress the importance of post-hoc evaluations in separate sections below. Our objective is not to be prescriptive about “best practices” although we do offer some insights where appropriate.

### Connectivity as conservation features

A simple and accessible way to integrate connectivity data into spatial planning is to treat it as a conservation feature, such that *r_ij_* is the amount of connectivity feature *j* (*e.g.* reproductive outflux) in planning unit *i*. In the classical minimum set problem (*i.e.* Marxan), a target is set for each feature, *T_j_*, (*e.g.* 50%) and the reserve system needs to contain at least that amount. This method can incorporate continuous or discrete conservation features (Figure 2). Examples of conservation targets best represented as continuous data include patch-level self-recruitment values (range between 0 and 1) or centrality measures (White *et al.* 2014; D’Aloia *et al.* 2017; Magris *et al.* 2018). In this treatment, a metric that represents the process of connectivity across the entire planning area receives a target. Definitions and potential conservation objectives for each connectivity based conservation feature available in Marxan Connect can be found on marxanconnect.ca.

**Figure 1:**
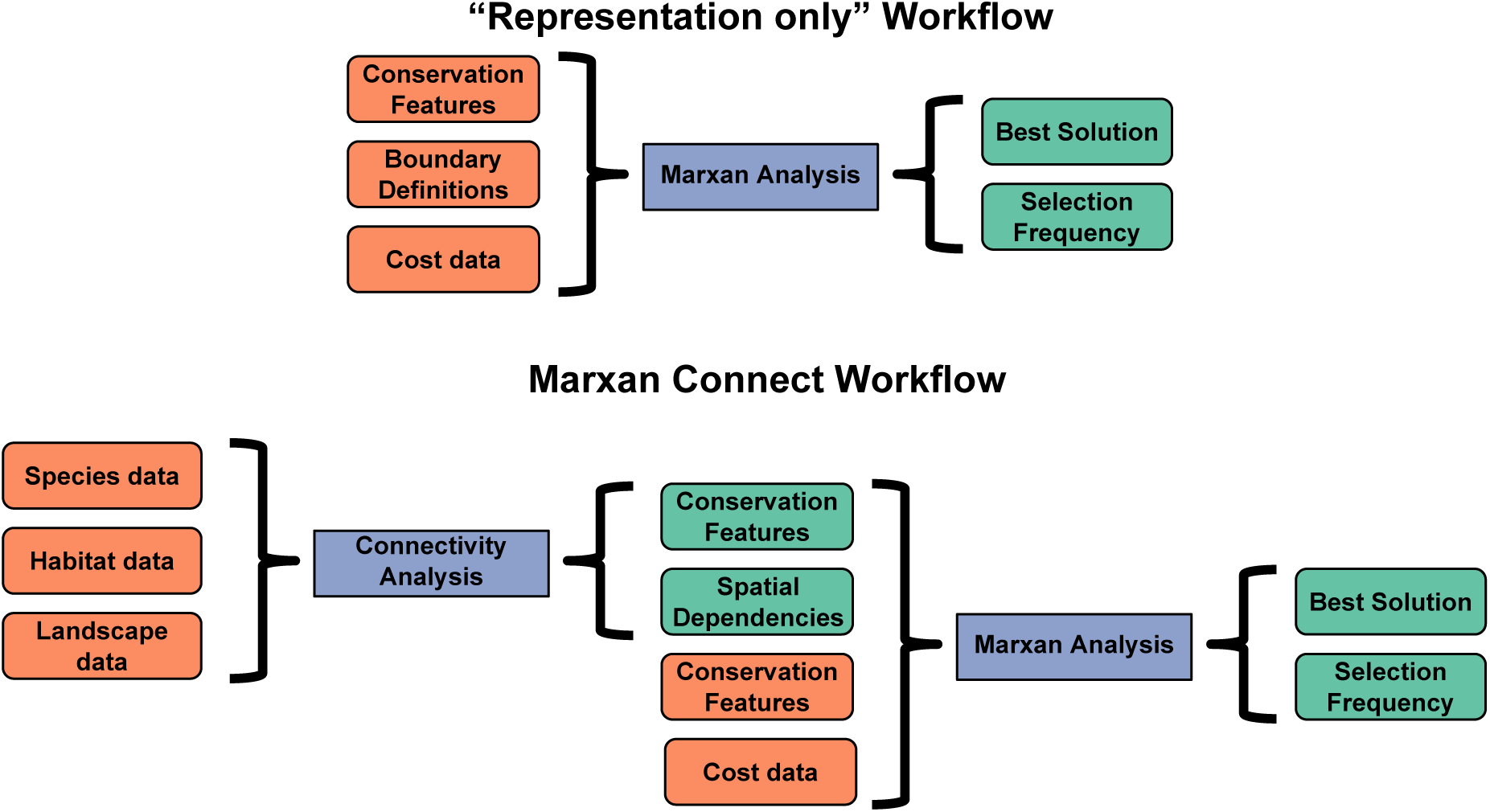
Comparison of workflows between the “representation only” approach to Marxan and “Marxan Connect.” Marxan Connect facilitates the use of connectivity data, derived from tagging data, genetics, dispersal models, resistance models, or geographic distance, by producing connectivity metrics and connectivity strengths (*i.e.* spatial dependencies that are used in the place of boundary definitions) before running Marxan. These connectivity metrics and linkage strengths are then used as inputs (connectivity-based conservation features or spatial dependencies) in the traditional Marxan workflow. The cost data in the traditional Marxan analysis refers to the cost of protecting a planning unit (*i.e.* opportunity cost), not the cost to traverse a landscape.

**Figure 2:**
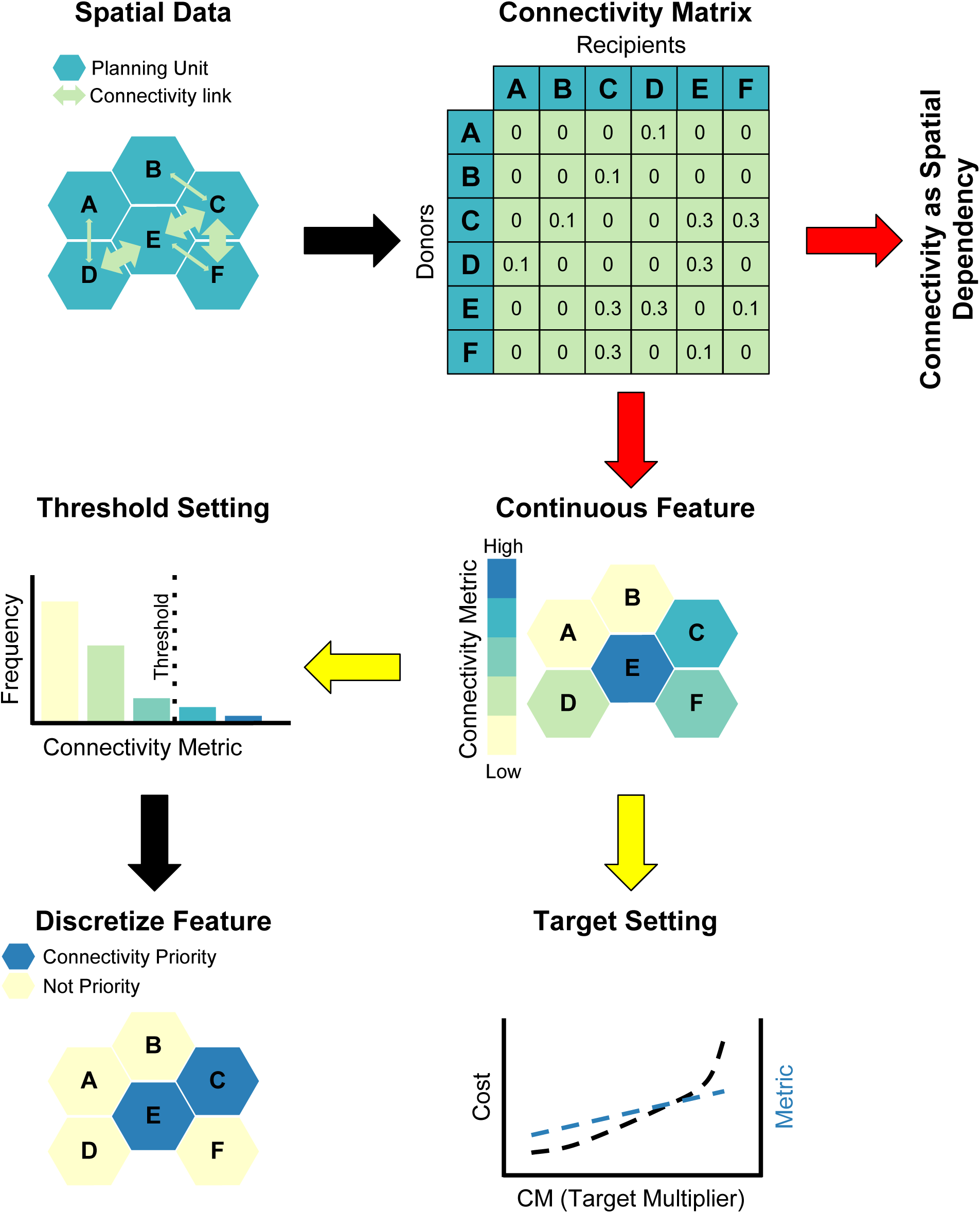
Data processing workflow for three possible methods for using raw connectivity data in spatial prioritization: 1) Connectivity as Spatial Dependency in the objective function (Raw Data ‐> Connectivity Matrix ‐> Connectivity as Spatial Dependency), 2) Continuous conservation features (Raw Data ‐> Connectivity Matrix ‐> Continuous Metrics ‐> Target Setting), or 3) Discrete conservation features (Raw Data ‐> Connectivity Matrix ‐> Continuous Metrics ‐> Threshold Setting ‐> Discrete Metrics. Coloured Arrows indicate important decision points. Red arrows indicate the conservation feature vs connectivity strength method decision point and yellow arrows indicate the continuous vs discrete conservation feature decision point

To increase the probability that the connectivity process is maintained, and that the spatial conservation plan is influenced by the conservation feature, a target higher than that of most other conservation features should be set. We suggest using a tunable ‘constraint’, a connectivity target multiplier (*C*), as a way to determine the higher target for connectivity-based conservation features (Figure 2 & 3). The target for the connectivity-based conservation feature, *T_c_*, would then be:

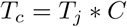

with *T_j_* being a typical conservation target for features not related to connectivity. An appropriate value for *C* can be determined by using a cost trade-off curve, similar to calibrating the BLM, where one would test the sensitivity of cost of the best solution and the total summed metric. It is worth noting that the BLM may interact with *C*. The value of *C* could be chosen as the divergent point, where the greatest increase in the connectivity metric is achieved for a relatively small increase in cost. However, the preferred approach would be to establish conservation targets leading to a reserve network design which meets ecologically relevant conservation objective(s), such as population viability or a within network metapopulation growth rate > 1 (See “Post-hoc evaluation” section for more details).

**Figure 3:**
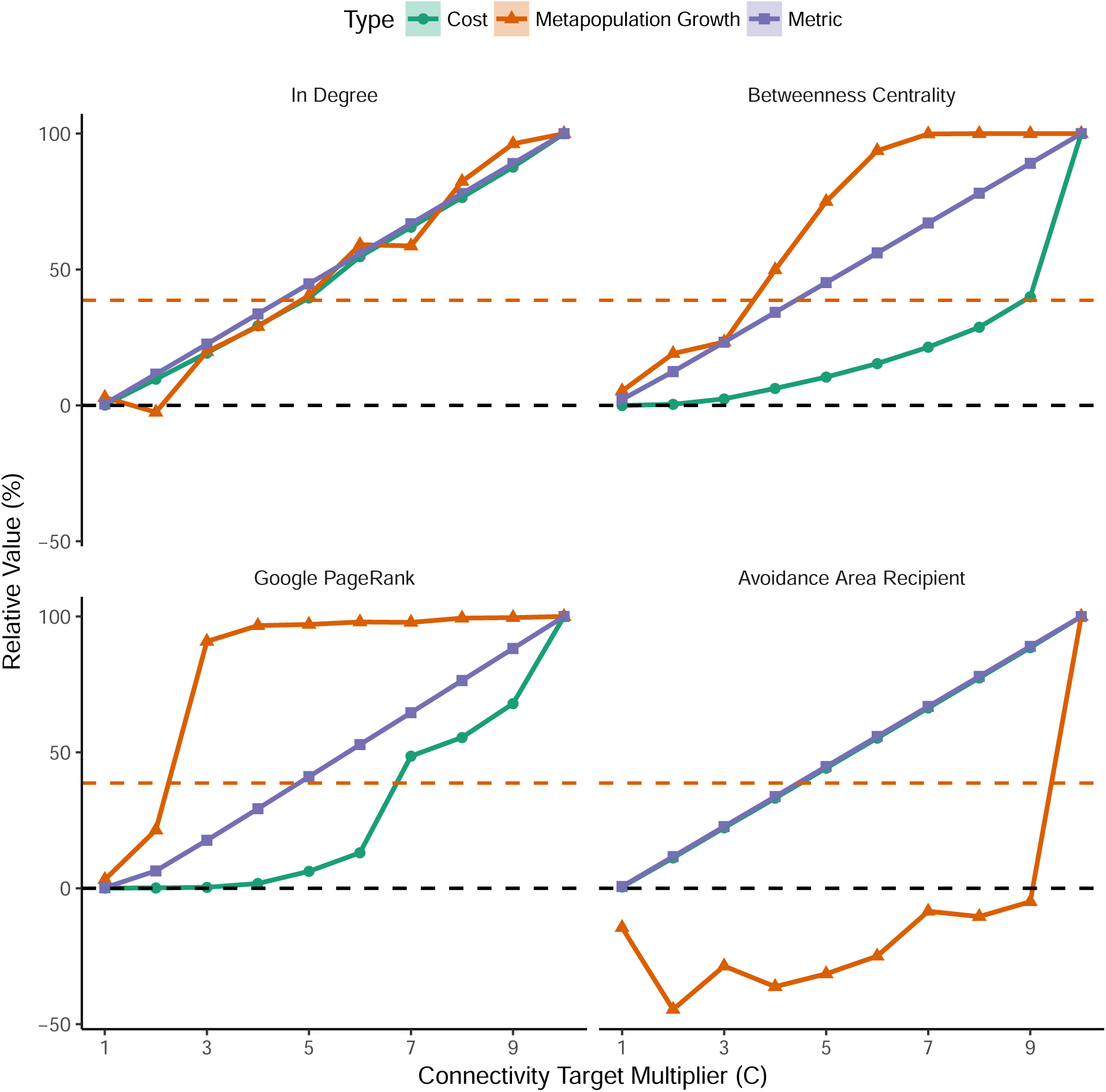
Connectivity target multiplier (C) sensitivity analysis, where the sum of the connectivity metric, estimated metapopulation growth rate (*λ_M_*), and cost of the selected network (mean ± S.E., n = 10) are plotted as relative percentages of their maximum values. Four connectivity metrics are used as examples: In Degree, Betweenness Centrality, Google Page Rank and Avoidance Area Recipient. The black dashed line is the value of the metric, growth rate, and cost of the selected network if the connectivity based conservation feature is not included. The orange dashed line indicates where *λ_M_* = 1, or the point at which the protected area network is self sustaining. The calculation of *λ_M_*, included here for demonstration purposes only, is estimated using the leading eigenvalue of a theoretical connectivity matrix which includes theoretical fecundity and survival (*i.e.* flow matrix). Full details on the generation of this figure can be found on marxanconnect.ca.

In contrast, conservation targets set for discretized connectivity conservation features *X_j_* can capture different levels of importance to connectivity amongst sites (Álvarez-Romero *et al.* 2018; Figure 2). To create a discrete feature, threshold value(s) must be chosen from the range of the continuous connectivity metric, ideally after ecological assessment or sensitivity analysis. The planning units that meet the threshold(s) are then discretized into unique features for which a target is set (Figure 2). Similarly, this type of threshold setting is often used with species distribution models, where each planning unit is assigned a probability of species occurrence, and a threshold value is used to convert these continuous distribution data to a binary map (presence vs absence or suitable vs unsuitable) to represent a particular species as a conservation feature (*e.g.* Minor *et al.* 2008). When appropriate, discrete and very high priority planning units could alternately be “locked-in” to the final solution to guarantee their inclusion. One disadvantage of setting a connectivity metric as a conservation feature is where the benefits of that connectivity conservation features are highly dependent on the final set of actions. For example, this approach currently does not allow the iterative recalculation of the metric as sites are added to the solution.

### Connectivity as spatial dependencies in the objective function

Another approach to incorporate connectivity in Marxan is using the directional connectivity-based data to inform the boundary cost (*cv_ih_*). This cost signifies the penalty associated with protecting one site but not other sites to which it is connected (Beger *et al.* 2010b). Here, a directional connectivity dataset is used to parameterize the boundary cost between all pairs of planning units (including non-adjacent units) and set the importance of incorporating connectivity with the BLM (or *b* in the equation in the “A spatial planning primer” section), also called “Connectivity Strength Modifier” (CSM) (Beger *et al.* 2010b). Increasing the BLM reduces the edge to area ratio by minimizing costs associated with unprotected adjacent boundaries; similarly, increasing the BLM in the connectivity context (*i.e.* CSM) will reduce the amount of leakage from the network (*i.e.* amount lost from the network) by minimizing costs associated with unprotected connectivity linkages, and thus maximising connectivity strengths across the entire Marxan solution.

Connectivity estimates as spatial dependencies in the objective function have been used to design marine protected area (MPA) networks in the Coral Triangle (Beger *et al.* 2010b, 2015). This approach can maximize the within-network connectivity, and may improve the metapopulation growth rate and other performance metrics. In Marxan Connect, one can combine the use of connectivity as spatial dependencies with a locked-in “Focus Area” (*e.g.* an existing protected area) to generate candidate stepping stones. However, the method will exclude isolated sites from the final solution unless these are included using other methods (*e.g.* a conservation feature for an isolated site which happens to contain a unique species).

### Connectivity as a Cost

A common approach to attributing costs in Marxan is to use inverse values as a treatment of the cost to be minimized. For example, such an approach might take the distance of a planning unit to the nearest port as a proxy of the cost to coastal fishing industries when establishing MPAs (Maina *et al.* 2015; Mazor *et al.* 2016; McGowan *et al.* 2017a). Here, the planning units closer to shore will be less costly than those farther offshore as the distance grows. Thus, an inverse distance cost makes the planning units closer to shore more expensive and less desirable for selection than those farther offshore. A more recently proposed method is to use connectivity as the cost to be minimized in the Marxan objective function. For example, Weeks (2017) used a “seascape connectivity cost” representing the inverse of the connectivity, expressed as the distance between adult habitat to nursery habitat. A disadvantage of the “connectivity as a cost” approach is that it precludes the consideration of other important socio-economic costs in the analysis, which are crucial for reducing conflicts with resource users and increasing the cost-effectiveness of implementation and management (Ban and Klein 2009). Further, this approach is not ideal since each planning unit’s contribution to the connectivity of the entire system relies on whether other sites are “in” or “out” of the reserve system.

### Connectivity-based objective function

Where the goal of including connectivity data into the spatial planning problem is to maximise the likelihood of the species’ persistence, then the most appropriate approach would be to include a persistence metric (*e.g.* population viability; metapopulation capacity) within the objective function. In this case, population connectivity together with fecundity, mortality and survival are included as a metapopulation model within the optimisation process. Currently, this is only realistic computationally for small problems (tens to hundreds of planning units) because the algorithm needs to calculate the performance metric very fast for simulated annealing to deliver good answers reasonably quickly. For example, Chollett *et al.* (2017) used a genetic algorithm to optimize an MPA network for maximum population persistence and fisheries yield, but this approach took 2 days on a high performance computing cluster which equates to ~300 days of single processor computing time. Even with powerful computational methods, such as integer linear programming (Beyer *et al.* 2016; Hanson *et al.* 2017), the problem formulation would be challenging as the performance metric of interest may vary non-linearly. Connectivity-based objective functions and their implementation in decision support software is a research priority.

## Making decisions: models, matrices and methods

Most data for the above methods focus on single species, yet many protected areas are designed to protect a diversity of species. There are many strategies for combining single-species connectivity data into a single multi-species connectivity matrix. Examples include taking the arithmetic or geometric means of multiple connectivity matrices, or connectivity metrics derived from them (Melià *et al.* 2016; D’Aloia *et al.* 2017). Others have calculated the probability of at least one species, or all species being connected (Jonsson, Nilsson Jacobi & Moksnes 2016; Magris *et al.* 2016). However, in all these cases, generating multi-species metrics or connectivity matrices resulted in some level of compromise that was suboptimal for a single species. If targeting conservation features, the most efficient solutions will be acheived by using single species conservation features that are representative of multiple life-stages and species with varying dispersal traits (Beger *et al.* 2015; Magris *et al.* 2016; Albert *et al.* 2017; D’Aloia *et al.* 2017). Alternatively, if the connectivity data are used to modify the boundary definitions, then a single connectivity dataset (*e.g.* edge list or matrix) per Marxan optimization must be used. Therefore, users are forced to calculate multi-species connectivity metrics, or run Marxan once per species and potentially combine the outputs. In all cases the consequences or trade-offs of the chosen strategy should be evaluated.

If the sole conservation objective is to maximize among-reserve connectivity, then modifying the boundary definitions and locking in existing reserves will likely produce the most efficient results. However, targeting connectivity features allows for more flexibility in objectives such as protecting areas that are important to maintain entire ecosystems (not just in reserves), avoiding areas with invasive species, or targetting areas with higher larval or adult spillover into unprotected sites. While these approaches may be slightly redundant, they are not mutually exclusive. Regardless of the data or method(s) that are chosen, post-hoc evaluations should be used to evaluate competing strategies.

The spatial and temporal scales of the connectivity data should be considered in all approaches. Ideally, connectivity should be quantified at the spatial scale of the planning units because the assumptions needed for rescaling can lead to erroneous results. The temporal scale of connectivity data should be aligned with the conservation objectives, such as providing demographic (*e.g.*, single generation movement) connectivity or safeguarding long-term gene flow (*e.g.*, many 100s of generations). Additionally, the planning area should extend beyond jurisdictional boundaries and the focus area to avoid edge effects which are particularly consequential for connectivity data (*e.g.* important source habitat may exist just beyond jurisdictional boundaries or focus areas).

## Post-hoc evaluation

None of the above methods, aside from customising the objective function, guarantee that population viability (see Tear *et al.* 2005) or metapopulation growth rate (*λ_M_* >; Figure 3) are maximised. This is because these methods are targeting the connectivity process per se but not the population outcomes explicitly. The use of simulated annealing for optimising complex performance metrics (*e.g.* population viability) currently has computational limitations. Therefore, the feasible solution to determine how well the final plan captures metapopulation outcomes in the analysis, or to compare the performance between plans, is to undertake post-hoc evaluations. The post-hoc analysis is structured as a sensitivity analysis, where multiple solutions are generated and compared to assess their performance in achieving the chosen connectivity objective. These solutions can be the result of varying Marxan parameters such as the conservation targets, the boundary length modifier, connectivity metrics, and/or costs, or the post-hoc analysis can compare individual spatial plans from the full ensemble created by the same input parameters and the simulated annealing process (Nicholson & Possingham 2006).

To illustrate this approach, we present a post-hoc sensitivity analysis to determine the optimal connectivity target multiplier value, *C*, across four connectivity metrics (Figure 3). This same approach could also be adapted for exploring the impact of using different methods, targets, thresholds, or data. We vary *C* in different Marxan scenarios using four different conservation features (in degree, betweenness centrality, Google PageRank and avoidance area recipients) to determine the impact on cost, and metapopulation growth. In this case, we used an in-reserve metapopulation growth rate greater than 1 (*e.g.* Figueira & Crowder 2006; Hale, Treml & Swearer 2015; marxanconnect.ca) as our ecologically relevant conservation objective. If the growth rate is lower than 1, the entire metapopulation would go extinct without external supplementation; however, this method requires detailed biological information. In this example, using “in degree” as a conservation feature increases the metapopulation growth rate linearly with cost, which has limited applicability. Both “betweenness centrality” and “Google PageRank” perform quite well with the latter being slightly better, likely because it considers the weight (*i.e.* strength) of the linkages. With “Google PageRank” as a conservation feature and *C* = 3 (*i.e.* conservation target of 30%), this example species is predicted to have a metapopulation growth rate > 1 and nearly the same level of growth as if the entire ecosystem was protected. In this example, “avoidance area recipient” performs extremely poorly because: 1) there was no parameter or mechanism in the population model that represented a reason to avoid the avoidance areas (*e.g.* impact of invasive species); and 2) the “avoidance area recipient” metric should always be discretized (*i.e.* low values of “avoidance area recipient” are desirable) since using the continuous metric would promote the selection of areas that receive the most propagules from the avoidance area.

In the second case, connectivity metrics can be calculated for individual conservation plans with varying spatial configurations that all meet the specified objective function (*i.e.* have the same input parameters). To identify the most dissimilar Marxan solutions in an analysis, a dissimilarity matrix (*i.e.* dendrogram) can be created using the Marxan cluster analysis function (Linke *et al.* 2011). The chosen metric can then be calculated for each plan to evaluate which spatial configuration best achieves the connectivity objective.

There are several different types of post-hoc assessments that can be performed, such as pattern-based assessments, evaluating metapopulation capacity, or using system models. In pattern-based assessments, for example, Krueck et al. (2017) developed metrics using local larval retention, import connectivity and export connectivity from connectivity matrices which can be calculated in a post-hoc analysis to evaluate how different conservation plans meet both conservation and fisheries objectives across protected and unprotected sites. Graph theoretic metric(s) or those that better capture population persistence can also be used, but few examples of their use in post-hoc analyses currently exist. In one example, Laita, Kotiaho, Monkkonen et al. (2011)explored how network connectivity measures (*i.e.* correlation length, expected cluster size, landscape coincidence probability, area-weighted flux, integral index of connectivity and probability of connectivity) changed with the addition of woodland key habitats to reserve networks in Finland. However, they highlight the need for a more detailed understanding of the caveats and justifications of these measures before they can be used for conservation purposes.

If additional demographic information of species, such as survival and mortality, is known, then the suggested course of action is to evaluate potential reserve networks using metapopulation models. With these models, it is possible to make predictions regarding the ecological outcomes such as the probability of going extinct in a certain time frame (Boyce 1992), the capacity to recover from a disturbance (Figueira & Crowder 2006), metapopulation lifetime (Kininmonth *et al.* 2010), probability of metapopulation extinction (Bode, Burrage & Possingham 2008), or other possible ecologically relevant conservation objectives. While increasing connectivity in reserve networks is generally desired; without models of metapopulation dynamics, connectivity risks becoming a relatively meaningless objective like “percent area covered by protected area” (Tear *et al.* 2005).

Lastly, system models are designed to simulate one or more processes related to the conservation objectives (*e.g.* prioritize stepping stones) or overall goals (*e.g.* population persistence). White *et al.* (2014) used population models to compare the performance of Marxan solutions generated with and without the inclusion of static larval connectivity information by calculating the equilibrium biomass (in and outside of protected areas) and fishery yield of the different spatial configurations in California. Similarly, tools such as the BESTMPA R package (Daigle, Monaco & Elgin 2017), allow users to test commercial fishery costs and benefits from various spatial conservation scenarios using a spatially explicit metapopulation model that interacts with fishing behaviour.

No matter what post-hoc analysis approach is used, selecting the most appropriate metric(s), understanding the caveats of the metric, and making ad hoc assumptions on how the user expects the metric to perform for the specified application is extremely important for interpreting and comparing the outcomes of these different measures (Pascual-Hortal & Saura 2006; see Laita, Kotiaho & Mönkkönen 2011). In conservation plans that incorporate existing protected areas, it is also important to evaluate the contribution of newly selected sites to the conservation objective, which can be accomplished by performing the post-hoc analysis with and without considering existing reserves (*i.e.* a gap analysis). This can reveal important information on the performance of the existing reserve system and can help ensure complementarity between the existing network and potential sites for protected area expansion.

## Marxan Connect

Because we recognize that there is considerable investment in Marxan-based prioritization, Marxan Connect was designed to help conservation practitioners incorporate connectivity into existing Marxan analyses. It guides users through:

1. Identifying and loading appropriate spatial data
  a. Planning units
  b. Focus areas
  c. Avoidance areas
2. Identifying and loading connectivity data
  a. Demographic-based
  b. Landscape-based
3. Calculates connectivity metrics or generates spatial dependencies
  a. Conservation features method
  b. Connectivity as spatial dependencies method
4. Optionally discretizes conservation features and exports Marxan files
5. Running Marxan
6. Evaluate results with basic plotting options

Marxan Connect allows users to export data products (*e.g.* connectivity metrics, Marxan files, etc.) at any of the above steps to enable users to base their workflow in or outside Marxan Connect.

For the landscape connectivity approach, Marxan Connect calculates connectivity metrics from networks based either on Euclidean distance or least-cost path between the centroid of planning units. However, other software packages such as Circuitscape (McRae, Shah & Mohapatra 2009) and Conefor (Saura & Torné 2009) currently provide a richer set of options and specialized methods. These software packages can be used to generate custom conservation features or connectivity matrices both of which can then be used in Marxan Connect. For example, one could generate a network using current density using Circuitscape and input the resulting connectivity matrix into Marxan Connect to generate conservation features or spatial dependencies.

For a user opting to use non-Marxan spatial prioritization software such as Zonation (Lehtomäki & Moilanen 2013) or prioritizr (Hanson *et al.* 2017), there is a high degree of compatibility with Marxan Connect. The approach of targeting conservation features is compatible with any spatial planning software. Certain software packages such as prioritizr can read Marxan-formatted files directly; therefore, Marxan Connect could be used with prioritizr to generate connectivity-related input files It also appears that modifying the boundary definitions with connectivity data could be performed with prioritizr’s “add_boundary_penalties” function. This is in addition to prioritizr’s “add_connected_constraints” function which tends to select unbroken chains of physically linked planning units (Önal & Briers 2006).

## Conclusions

The approaches for including connectivity in spatial planning are rapidly evolving and few “best practices” exist. Here, we provide some guidance on methods, data sources, and models, as well as a novel open-source tool to support these methods. However, connectivity-based conservation targets are ecologically meaningless unless placed in the context of broader ecologically relevant conservation objectives such as population viability, expected time to extinction, or metapopulation growth rate. Similarly, connectivity is usually only one criterion in planning, and will be considered alongside area-based targets, socio-economic goals, and multi-species requirements.

Connectivity is a complex topic with abundant terminology and a diversity of methods that require substantial effort to understand and apply to spatial prioritization scenarios correctly (Beger et al. in prep). If connectivity is to widely inform protected area planning, communication channels between experts in the fields of connectivity and population dynamics and planners must be improved. The experts, in particular, should make their research outcomes more accessible to practitioners by providing openly available data and clarifying definitions, assumptions, and limitations. For example, the term “connectivity matrix”, while central to the concept of connectivity, does not provide enough information to spatial planners or even to other connectivity experts to incorporate connectivity into spatial planning initiatives. With Marxan Connect, we hope to offer standardized methods and terminology to help close this research-implementation gap.

While Marxan Connect represents an advance in facilitating the incorporation of connectivity into the design of protected areas, it does not guarantee that reserves will be “well connected”. Only post-hoc evaluation of the reserve design related to ecologically relevant conservation objective(s) can inform practitioners of the resilience and persistence of targeted populations. However, the tools provided in Marxan Connect greatly improve the likelihood that a selected reserve design will adequately meet those conservation objective(s).

## Acknowledgements

Funding for the development of this software was provided by the Natural Sciences and Engineering Research Council of Canada through the Canadian Healthy Oceans Network (NSERC NETGP 468437-14), and the Discovery Grants program (NSERC DG 34851-2012), and the University of Queensland. Additional contributions were provided by the University of Leeds, The Nature Conservancy, Dalhousie University, The University of Melbourne, and the Australian Research Council - Centre of Excellence for Environmental Decisions (CEED). This project builds on the existing Marxan (Ball, Possingham & Watts 2009) software and would not be possible without the hard work of Ian Ball, Matt Watts, and Hugh Possingham. The authors also wish to thank Ryan Stanley, Marco Andrello, and Jo Clarke for constructive feedback on early versions of the software or manuscript.

## Author’s contributions

AM and MB conceived the application and acquired funding; RD developed the application and website; RD, AM, AB, and MB worked on the initial development and early testing of the application. All authors made significant contributions to the later development stages of the application and website. All authors contributed critically to making improvements to the application, drafting the manuscript, and provided final approval for publication.

## Supplementary Material

The source code for the software and website can be found at https://github.com/remi-daigle/MarxanConnect. There are repeated references in the tutorial and glossary section of marxanconnect.ca, the website may evolve as the software is improved. The original publication version of the website and app have been archived on Zenodo (Daigle et al. 2018 *This will be archived and added to references upon acceptance of the manuscript*)

